# Influence of transglutaminase mediated crosslinking on the structure-function-digestion properties of *Lupinus angustifolius* protein evaluated using a multiscale approach

**DOI:** 10.64898/2026.03.18.712645

**Authors:** A. Mukherjee, D. Duijsens, I. Fraeye, F. Weiland, T. Grauwet, I. Van de Voorde

## Abstract

This study presents a multidisciplinary approach to evaluate the structure formation and digestion of lupin protein crosslinked with transglutaminase (TG). TG was applied at 0–10 U/g protein, and structural development was assessed by oscillatory rheology (G’, G”), while SDS-PAGE and o-phthaldialdehyde (OPA) assays were used to evaluate protein participation and the reduction of free ε-amino groups, respectively. Proteomics was further employed to characterise molecular features associated with crosslinking behaviour. Lupin protein showed a clear dose-dependent increase in gel strength during incubation, with G’ values reaching 214 ± 43.9 Pa at 10 U/g TG, compared to 7.2 ± 0.6 Pa in the untreated control. Across all conditions, G’ remained higher than G” throughout frequency sweeps, and low tan δ values confirmed the formation of elastic networks driven by covalent crosslinks. SDS-PAGE and OPA results consistently demonstrated efficient crosslink formation, which increased with both incubation time and TG dosage, with SDS-PAGE indicating involvement of specific protein fractions. Proteomic analysis revealed disordered structural domains in the protein are preferred regions to form crosslinks. Furthermore, TG treatment was found to slow the digestibility of the crosslinked lupin protein. Overall, this work demonstrates how integrating proteomic insights with functional measurements can guide the selection and optimisation of plant proteins for enzymatic structuring. The approach offers a rational pathway to enhance the functionality of alternative protein sources such as lupin, supporting the development of sustainable food systems, including applications in meat and dairy analogues.

## 1 Introduction

The growing demand for sustainable, plant-based protein sources has intensified interest in legume proteins as alternatives to animal-based counterparts. Among legumes, lupin (*Lupinus* spp.) has emerged as a promising alternative to soy due to its high protein content (30–45% of seed weight) and ability to grow in temperate environments. Lupin protein also has a favourable amino acid composition (low in sulphur containing amino acids and high in lysine) to be complemented well with cereals (high sulphur containing amino acids, low in lysine) (Gulisano et al., 2019; Shrestha et al., 2021). According to the European Union plant database, *Lupinus angustifolius* (narrow-leafed lupin) accounts for most commercial lupin varieties in Europe (European Commission, 2024).

Despite their potential, lupin proteins can exhibit limited functionality, particularly with respect to gelation, which is critical for their incorporation into plant-based foods such as cheese and meat alternatives (Al-Ali et al., 2021; Chew et al., 2003). Batista et al. (2005) and Berghout et al. (2015a) reported lupin protein to have the poorest gelling ability compared to soy and pea proteins, even when tested at a higher protein concentration. Lupin protein has more disulfide bonds than soybean protein, making it highly thermally stable. This stability limits protein unfolding during heating, reducing its ability to form a strong gel network (Duranti et al., 2008). In contrast, soybean protein, with fewer S–S bonds, unfolds more readily and forms stronger gels thermally (Sirtori et al., 2010). Although lupin protein contains free sulfhydryl groups, the protein remains thermally stable and compact, hence those SH groups do not efficiently participate in forming new intermolecular disulfide bonds (Berghout et al., 2015b). One strategy to improve functional performance is cold-set gelation via enzymatic crosslinking, particularly using microbial transglutaminase (TG) (Isaschar-Ovdat & Fishman, 2018; Maltais et al., 2008). TG catalyses an acyl-transfer reaction between the γ-carboxamide group of glutamine residues and the ε-amino group of lysine residues, forming covalent ε-(γ-glutamyl)-lysine isopeptide bonds (Tatsukawa et al., 2020). This crosslinking modifies protein structure and intermolecular interactions, possibly leading to enhanced gel strength, water-holding capacity, and stability in food systems (Chen et al., 2018; Qin et al., 2016).

TG has been extensively studied in legume systems, such as soy, pea and mung bean proteins, where it promotes the formation of high-molecular-weight aggregates and significantly improves textural properties (Djoullah et al., 2018; Mukherjee et al., 2026; Schlangen et al., 2023). Santoso et al. (2024) crosslinked a 1:1 blue lupin and whey protein dispersion with 13 U TG/g protein. SDS-PAGE analysis indicated extensive TG-induced crosslinking, particularly involving lupin proteins, enhancing thermal and emulsion stability. Ceresino et al. (2021) explored how TG-mediated crosslinking combined with lipidic additives (glycerol, lecithin, linoleic acid) could systematically modulate the *micro-and nano-architecture* of lupin protein-based frozen-cast foams. However, the efficiency of TG-mediated crosslinking is largely governed by the molecular characteristics of the target proteins, particularly the accessibility of reactive residues such as lysine and glutamine (Kang et al., 2024). In this context, the structural heterogeneity of lupin proteins becomes highly relevant. Lupin contains four major storage protein families, collectively termed conglutins: α-conglutins (11S legumin-like globulins), β-conglutins (7S vicilin-like globulins), γ-conglutins (7S basic globulins), and δ-conglutins (2S albumin-like proteins) (Nadal et al., 2011). α-and β-conglutins are the predominant fractions, together accounting for 80-90 % of the total seed protein, followed by γ-conglutin (15–20 %) and δ-conglutins (<10 %) (Náthia-Neves et al., 2025). Their structural diversity and solubility characteristics influence both the techno-functional and digestion properties of lupin protein ingredients (Dhakal et al., 2024). Lupin proteins differ from other legume proteins in both molecular composition and tertiary structure. For instance, β-conglutins are particularly hydrophilic and structurally flexible compared with vicilins from other legumes. α-conglutins resemble more legumin-like structures and have distinct acidic and basic chains that may vary in accessibility to TG (Duranti et al., 2008). These differences may lead to selective preferences for crosslinking, influencing the formation of macroscopic gels. Rheological analyses to study the viscoelastic properties of the formed gels are crucial to understand TG-mediated crosslinking of lupin protein but are currently still very limited. Existing studies report only basic flow behaviour or structural outcomes (Ionescu et al., 2009) rather than a detailed viscoelastic characterisation of gel networks (storage and loss moduli) that would directly elucidate TG-mediated crosslinking effects on structure formation. Moreover, examining the effects of TG-mediated crosslinking at the molecular level is essential to better interpret the observed structural and functional changes. Proteomic approaches are very promising to study how TG interacts with specific lupin proteins, which is critical for optimising functional outcomes and tailoring ingredient properties. A previous study by Mukherjee et al. (2026) revealed preferential crosslinking of convicilin and vicilin proteins of pea protein by TG using proteome analyses. Schäfer et al. (2005) crosslinked 13 % blue lupin protein with 0.1 g TG/g protein. They quantified ε-(γ-glutamyl)lysine dipeptide formation via HPLC-MS analysis in TG crosslinked lupin protein, but an in-depth molecular understanding is currently lacking. Moreover, most studies use a single TG dosage, thus overlooking the influence of the protein-enzyme ratio on structure formation while TG dosages are often reported in mass units rather than enzymatic activity, which complicates comparison across studies and limits interpretation of treatment effects.

Beyond functionality, TG-mediated crosslinking can also have nutritional implications, especially regarding protein digestibility (Glusac et al., 2020; Romano et al., 2016a). The formation of covalent isopeptide bonds may hinder enzymatic hydrolysis during gastrointestinal digestion, potentially reducing the bioavailability of amino acids or altering the release of bioactive peptides (Feng et al., 2024). While some studies report moderate reductions in *in vitro* digestibility following TG treatment of legume proteins, the effect is highly dependent on the structural organisation of the network and the degree of crosslinking (Tang et al., 2008; Wang, Xin, et al., 2022a). Furthermore evaluating digestive fate of structured protein is important for understanding how processing and structure impact digestion in order to design foods with targeted nutrient release (Verkempinck et al., 2022). Given the increasing consumer demand for nutritionally balanced plant-based foods, it is essential to understand how TG-mediated structural changes impact the digestibility of lupin proteins.

In conclusion, it is clear that despite lupin protein has important structuring potential, impact on TG-mediated crosslinking on lupin protein remains poorly understood. Existing studies are limited, fragmented, and primarily confined to bulk analytical techniques such as SDS PAGE, offering only minimal molecular insights, and providing little to no understanding of the digestion properties of crosslinked lupin proteins. To address these knowledge gaps, this study investigates the effect of TG-mediated crosslinking on the structural and nutritional properties of lupin protein. Using small amplitude oscillatory shear rheology, gelation was monitored as a function of TG dosage and incubation time. To elucidate the molecular basis of crosslinking, SDS-PAGE and advanced proteomics were employed to identify which protein domains specifically affect crosslinking efficiency. Finally, implications of the crosslinking on protein digestion kinetics was assessed in order to link crosslink density and subunit-level modifications to protein digestion kinetics.

## 2. Materials and methods

### 2.1. Materials

BP80F *Lupinus angustifolius* protein isolate was received from Wide Open Ingredients (Leederville, WA). The ingredient was selected to reflect industrially relevant material characteristics, as commercial protein isolates represent the predominant form used in food applications. A Ca^2+^-independent TG enzyme was acquired from Novozymes (Galaya® Prime, Denmark). The activity of the applied enzyme batch was determined according to the method of Djoullah et al. (2015a) and was found to be 116.8 U/g enzyme. All chemicals used were of analytical grade. LC grade chemicals were used for proteome analyses. Enzymes and reagents for simulated digestion were provided by Sigma Aldrich (Belgium).

### 2.2. Lupin protein isolate characterisation

#### 2.2.1. Proximate composition

The protein content of the lupin protein isolate was determined using the Kjeldahl method (ISO 20483) with a nitrogen to protein conversion factor of 6.25 (Roman et al., 2025). The lipids were extracted using chloroform/methanol (1:1) followed by a gravimetric determination of the total lipid content (Ryckebosch et al., 2012). The moisture content of the protein isolate was assessed by the oven drying method (ISO 24457) and total mineral content was estimated by dry ashing (ISO 2171). All analyses were done in triplicate.

#### 2.2.2. Differential scanning calorimetry

Thermal denaturation properties of the protein in the isolate were studied based on the method of (Chigwedere et al., 2018). A differential scanning calorimeter (DSC Q2500, connected with a refrigerated cooling system RCS40, TA instruments, USA) was applied for quantitative determination of the denaturation enthalpy and transition temperatures. Briefly, 10 mg of the sample was placed in a high-volume Tzero pan (TA instruments, USA), and demineralised water was added at a 1:3 (w/v) ratio. A temperature ramp was applied from 20 to 120 °C at 5 °C/min with a sampling interval of 0.10 s/point. The DSC chamber was flushed with nitrogen gas at a flow rate of 50 mL/min. Trios software (v5.7.2.101, TA Instruments, USA) was used to analyse the obtained thermograms. The analysis was conducted in triplicate.

#### 2.2.3. Solubility

Protein solubility was determined in triplicate based on previous work (Mukherjee, Van Pee, et al., 2026). The lupin protein isolate was dispersed in a phosphate buffer (50 mM sodium phosphate containing 0.3 M NaCl, pH 7) in a concentration of 1% (w/w) and stirred on a magnetic stirrer for 1 h at room temperature. Subsequently, the dispersion was centrifuged for 20 minutes at 20 °C at 8960 g (Centrifuge 5910i, Eppendorf, Germany). The protein content of the collected supernatant was analysed using the Kjeldahl method (see section 2.2.1). Protein solubility was calculated as follows:

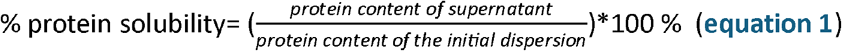

### 2.3. Enzymatic crosslinking of lupin protein with TG

Enzymatic crosslinking of the lupin protein isolate was conducted following the method described for pea protein in Mukherjee et al. (2026). At first, 0.5 g of lupin protein was added to a reaction tube and diluted to 5.0 g with phosphate buffer (50 mM sodium phosphate containing 0.3 M NaCl, pH 7) to reach a 10 % (w/w) protein dispersion. This protein concentration was selected based on preliminary experiments and previous work on pea protein (Mukherjee, Van Pee, et al., 2026). Dispersions were stirred for 1 h at room temperature on a magnetic stirrer. After that, a defined amount of TG was added to the reaction tubes according to the intended enzyme dosage (0.25, 0.5, 1, 5, 10 U TG/g protein). The reaction tubes were then incubated up to five hours at 40 °C in a water bath. Subsequently, each reaction tube was subjected to a thermal treatment at 70 °C for 10 min to inactivate the enzyme. Blank samples were also included, which underwent one of the following two treatments: i) **n**o **e**nzyme addition, no thermal incubation, and **n**o **h**eat inactivation treatment (hereafter referred to as NHNE); or ii) **n**o **e**nzyme addition and no thermal incubation, but **h**eating under inactivation conditions (hereafter referred to as HNE). After that, samples were freeze-dried and kept at -20 °C until further analysis.

### 2.4. Determination of structure formation using rheology

#### 2.4.1. Dynamic oscillatory rheological measurements

The viscoelastic properties of lupin protein isolate crosslinked with TG were assessed using small amplitude oscillatory shear rheology according to Mukherjee et al. (2026) to investigate structure formation. A stress-controlled rheometer (AR 2000ex, TA Instruments, New Castle, USA) equipped with cross-hatched parallel plate geometry (40 mm diameter) was used for the analyses. The gap was set at 1 mm. Protein dispersions were prepared and varying TG dosages were added as described in section 2.3, including a control (0 U TG/g protein). After brief vortexing, the dispersions were immediately loaded onto the rheometer at 20 °C. A solvent trap with a covering hood was used to limit evaporations. Throughout the analyses, the storage modulus (G’), loss modulus (G”), and tan δ (G”/G’) were recorded. All analyses were done in duplicates.

After loading the sample, the plate temperature was increased to 40 °C at a rate of 5 °C/min. A time sweep was then conducted over a period of 5 hours to monitor structure formation as a function of incubation time. Following this, the temperature was raised to 70 °C at 5 °C/min and held for 10 minutes at 70 °C to inactivate the enzyme. The sample was then cooled to 20 °C at 5 °C/min and held for 30 minutes at 20 °C. During the entire time sweep, a constant strain of 0.5 % and frequency of 1 rad/sec were applied.

Frequency sweep measurements were carried out on the gels formed immediately following the above-described steps. These measurements spanned a frequency range of 0.062 to 62 rad/sec, with 20 measuring points per decade. The strain was held constant at 0.5 %, and the temperature was maintained at 20 °C.

To determine the linear viscoelastic region (LVR) of the gels, a strain sweep was performed. Samples were subjected to strains ranging from 0.0001 to 10 at a constant temperature of 20 °C and a frequency of 1 rad/sec with 20 measuring points per decade. The LVR was established by identifying the oscillatory strain at which G’ deviated by more than 10 % from the average G’ of the preceding five measurements.

#### 2.4.2. Data analysis

The frequency sweep data were fitted using a power-law model (Ikeda & Foegeding, 1999; Mukherjee, Van Pee, et al., 2026):

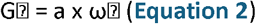

where a is the consistency coefficient related to the gel strength (Pa-s⍰) and b is the frequency exponent describing the dependence of G⍰ on angular frequency (ω, rad-s^−1^). Parameter estimates were obtained from duplicate measurements. Differences in mean rheological parameters obtained from time sweep and strain sweep tests were assessed by one-way analysis of variance (ANOVA), followed by Tukey’s HSD post hoc test at a significance level of 0.05. All statistical analyses were conducted using JMP Pro 17 (SAS Institute Inc., Cary, NC, USA).

### 2.5. Determination of free ε-amino group content

#### 2.5.1. Spectrophotometric analysis

Measurement of the reduction in free ε-amino groups resulting from TG-mediated crosslinking between lysine and glutamine residues can be utilised to indicate the progress of crosslinking. The free ε-amino groups in the samples were quantified using o-phthaldialdehyde (OPA) as mentioned in (Mukherjee, Van Pee, et al., 2026). The crosslinked protein samples, together with NHNE and HNE samples (50 mg; see Section 2.3), were suspended in 1 mL of 0.1 M sodium phosphate buffer containing 2% (w/v) SDS at pH 8 and vortexed for 30 s. The resulting mixture was centrifuged at 5641 × g for 10 min at 20 °C using a Centrifuge 5424 R (Eppendorf, Germany). For spectrophotometric analysis, 3 mL of freshly prepared OPA reagent was added to 400 µL of the diluted (in 0.1 M sodium phosphate buffer, pH 8) supernatant, followed by incubation in the dark for 2 minutes at room temperature. After incubation, absorbance was measured at 340 nm using a spectrophotometer (Cary 100 UV-Vis, Agilent Technologies, USA). A standard curve of L-serine (25–150 mg/L) was used to convert the absorbance values into L-serine equivalents and subsequently into the concentration of free ε-amino groups.

#### 2.5.2. Data analysis

SAS version 9.4 (SAS Institute, Inc., Cary, NC, USA) was used for kinetic modelling of the decline in free ε-amino groups. A fractional conversion model (Verkempinck et al., 2024) was used to simulate the data following equation:

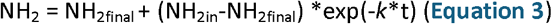

where *k* is the estimated rate constant of crosslinking (min^-1^), t is the incubation duration (min), *NH*_*2final*_ is the final content of ε-amino groups (mg/g sample), and NH_2in_ is the initial content of ε-amino groups (mg/g sample). The measurements can be regarded as independent observations of the system at different time periods because each time point in the crosslinking study was evaluated using a separate reaction tube, allowing for kinetic modelling of the data (Duijsens et al., 2022).

### 2.6. SDS-PAGE

SDS-PAGE (Laemmli,1970) was employed to identify the involvement of specific protein subunits of lupin samples in the crosslinking process. The method was based on previous work by Mukherjee, Van Pee, et al. (2026). Protein dispersions (0.5 % (w/v)) in 50 mM sodium phosphate buffer containing 0.3 M NaCl, pH 7) using the freeze-dried NHNE, HNE and the crosslinked samples after incubation for 0, 30, 60, 120, 180, 240, and 300 minutes (see section 2.3) were prepared, followed by vortexing for 1 minute. The dispersions were diluted 1:1 in sample buffer containing 95% Laemmli Sample Buffer (Bio-Rad Lab., Hercules, CA, USA) and 5 % β-mercaptoethanol (Bio-Rad Lab., Hercules, CA, USA) and then heated at 100 °C for 4 minutes. After cooling to room temperature, samples (30 µl) were loaded onto a gel (Bio-Rad, Criterion TGX, Any kD™, Stain-Free) along with protein standards (15 µl, Precision Plus Protein Unstained, Bio-Rad). The electrophoresis chamber was filled with 800 mL of Tris/Glycine/SDS buffer (Bio-Rad Lab., Hercules, CA, USA). Protein separation was performed at a constant current of 20 mA for 25 minutes, followed by a constant voltage of 300 V for 20 minutes. The protein bands were then visualized using the ChemiDoc Mp Imaging System (Bio-Rad Lab., Hercules, CA, USA).

### 2.7. Proteome analysis

#### 2.7.1. Hydrolysis of crosslinked proteins

The tryptic hydrolysis of the (non)crosslinked lupin protein was conducted based on prior work (Mukherjee, Van Pee, et al., 2026).The lyophilised crosslinked protein sample (see section 2.3), incubated with 10 U TG/g protein for 5 h (n = 4), along with the NHNE sample (n = 4), were resuspended at a concentration of 10 mg/mL in buffer containing 8 M urea (Merck KGaA, Darmstadt, Germany) and 50 mM ammonium bicarbonate (Sigma-Aldrich), pH 8.5. Protein solubilisation was achieved using an ultrasonication probe (two pulses of 30 s at 50 % amplitude, 120 W, with a 1 min interval) (Thermo Scientific, Waltham, MA, USA). Samples were kept on ice during sonication to minimise heating. Protein concentration was determined using the bicinchoninic acid (BCA) assay (Pierce™ BCA Protein Assay Kit, Thermo Scientific, Waltham, MA, USA). An aliquot containing 50 µg protein was reduced and alkylated at room temperature for 45 min with 10 mM tris(2-carboxyethyl)phosphine and 25 mM chloroacetamide (both Sigma-Aldrich). Proteolytic digestion was performed by adding 1 µg Trypsin/LysC (Thermo Fisher Scientific, Waltham, MA, USA) and incubating overnight at 37 °C. Peptides were subsequently desalted using in-house packed C18 StageTips (Empore SPE Disks, 3M, Saint Paul, MN, USA) and freeze-dried.

#### 2.7.2. Tandem mass tag (TMT) labelling

TMT Pro 16-plex reagents (Thermo Scientific, Waltham, MA, USA) were used to conduct isobaric labelling of the peptides as described by Mukherjee et al. (2026) with modifications. Briefly, 0.5 mg of each TMT reagent was dissolved in 32 µL of anhydrous acetonitrile (Sigma-Aldrich) following the manufacturer’s instructions. Freeze-dried peptides (section 2.7.1.) were reconstituted in 50 mM 2-[4-(2-hydroxyethyl) piperazin-1-yl] ethanesulfonic acid (HEPES; Sigma-Aldrich), pH 8.5 (Zecha et al., 2019), and 6 µL of the appropriate TMT reagent was added. Labelling was performed for 2 h at room temperature. Then the labelling reaction was quenched by adding hydroxylamine (Sigma-Aldrich) to a final concentration of 0.5 % (w/v). The labelled peptides were then combined, desalted using in-house packed C18 tip columns (Empore SPE Disks, 3M, Saint Paul, MN, USA), and freeze-dried.

#### 2.7.3. Liquid chromatography -tandem mass spectrometry (LC-MS/MS) analysis

Peptide digests were analysed using an Orbitrap Astral mass spectrometer (Thermo Scientific, Waltham, MA, USA) interfaced with a Vanquish Neo UHPLC system (Thermo Scientific). The method was based on previous work (Mukherjee, Duijsens, et al., 2026) Chromatographic separation was performed using mobile phases consisting of buffer A (0.1 % formic acid) and buffer B (80 % acetonitrile containing 0.1 % formic acid) (Sigma-Aldrich). A gradient optimised for a 60 samples-per-day workflow was applied, with peptides initially retained on a PepMap Neo C18 trap cartridge (Thermo Scientific, Waltham, MA, USA) and subsequently separated on an Aurora Ultimate column (IonOpticks). Peptides were eluted at a flow rate of 500 nL/min using a gradient that increased from 1 % to 8 % buffer B over 0.8 min, followed by a rise from 8 % to 38 % buffer B over 15.5 min, and a final increase from 38 % to 50 % buffer B over 1.3 min. The column was then washed and re-equilibrated prior to the next injection. Mass spectrometric analysis was carried out in positive ion mode using an EasySpray ion source, with the spray voltage set to 2.0 kV and the ion transfer tube temperature maintained at 290 °C. Data were acquired in data-dependent acquisition (DDA) mode, with each cycle consisting of a full MS (MS1) scan followed by MS/MS (MS2) scans. MS1 spectra were acquired in profile mode at a resolution of 180,000 over an m/z range of 380–1500, using a custom automatic gain control (AGC) target of 300 %, a maximum injection time of 25 ms, and a single microscan. Monoisotopic peak selection was enabled for peptide ions with charge states between 2 and 6. The acquisition was set to cycle-time mode, with a 0.5 s interval between master scans and a dynamic exclusion duration of 20 s. MS2 spectra were collected in centroid mode using a 0.4 m/z isolation window and higher-energy collisional dissociation (HCD) at 32 % collision energy. A custom AGC target of 500, a maximum injection time of 20 ms, and one microscan were used. Fragment ions were detected using the Astral analyser with TMT mode enabled over a scan range of 110–2000 m/z.

#### 2.7.4. Data processing and statistical analysis

The mass spectrometry raw data was analysed as described in Mukherjee et al. (2026), deviating in the FASTA file used. The FASTA was a combined *Lupinus angustifolius* and *Helianthus annuuss* file with 119,652 entries, downloaded from Uniprot on the 23^rd^ February 2026. The inclusion of *H. annuus* sequences was required due to the experimental design, in which samples from both species were analysed within the same TMT multiplex experiment. Distinct TMT labels were assigned to each species (e.g., *L. angustifolius* samples labeled with TMT126–TMT130N; eight labels in total), but the combined LC–MS/MS acquisition required searching against a database containing proteins from both species to ensure correct peptide assignment and to prevent cross-species misidentification. Statistical analysis was conducted as described in Mukherjee et al. (2026), using the same scripts with adjustments. In brief, the raw TMT reporter intensities were transformed and calibrated using variance stabilising normalisation (Huber et al., 2002) and statistical testing was conducted using limma (Ritchie et al., 2015). Testing for enrichment of cross-linking sites within disordered regions was performed using a Fisher’s exact test. All analyses were conducted using R (version 4.5.1) in the RStudio environment (version 2025.09.2 Build 418) using following libraries: stringr (Wickham, 2023), dplyr (Wickham, François, et al., 2023), ggplot2, vsn (Huber et al., 2002), limma (Ritchie et al., 2015), ggrepel (Slowikowski, 2024), ggpointdensity (Kremer, 2019), scales (Wickham, Pedersen, et al., 2023), ggseqlogo (Wagih, 2024), rmotifx (Wagih et al., 2016), and viridisLite (Garnier et al., 2023). The mass spectrometry raw data can be downloaded from ProteomeXchange (Deutsch et al., 2023) or jPOSTrepo (Okuda et al., 2017) with the identifier PXD075138 / JPST004447 (the data analysis scripts are available from Zenodo via https://doi.org/10.5281/zenodo.18846970.

For the calculation of the crosslink heuristic, the complete Uniprot database including amino acid sequences and protein domain information was downloaded on 12^th^ January 2024 (251,702,059 protein entries from 1,311,639 organisms) and split into chunks of 2,000,000 lines. The corresponding Gene Ontology (GO) terms for each protein were downloaded from Uniprot on 29^th^ April 2024. For each organism, the number of proteins, the number of disordered protein regions within a protein (in bins of 0, 1, 2 and ≥3), the number of lysines (K) and glutamines (Q) within these regions as well the number of these K with a glutamic acid (E) at position –7 and Q with an E at position –3 was enumerated (Mukherjee et al., 2026). The (arbitrary) heuristic for each organism was calculated as the percentage of multiconnector (≥ 3 K or Q within disordered regions) and chain-former (2 K or KQ within disordered regions) storage proteins (GO:0045735) within all storage proteins. This was done to introduce a weight for protein abundance within the organisms, with storage proteins expected to be the most abundant ones. This percentage was then multiplied by the balance of K and Q within disordered regions, the same number of K and Q being preferred as TG cross-links between them. Afterwards the list was ranked according to the heuristic, with higher values expressing a higher percentage of multi-connectors/chain-formers within storage proteins and/or a more equal number of K and Q within disordered regions. Afterwards, the list was filtered for entries of the class Magnoliopsida and organisms with < 1000 protein entries were excluded to remove potentially incompletely sequenced entries. Analysis scripts are available via Zenodo under the URL https://doi.org/10.5281/zenodo.18886960.

### 2.8. In vitro digestibility assessment of the (crosslinked) lupin protein

#### 2.8.1 INFOGEST static in vitro digestion

The effect of TG-mediated crosslinking on lupin protein digestibility was assessed using the INFOGEST static *in vitro* digestion protocol (Brodkorb et al., 2019). Samples with the clearest differences in crosslinking (0 and 10 U TG/g protein) were selected for the digestibility study. The crosslinked lupin protein was prepared as mentioned in section 2.3. Incubation with TG was conducted for 60 min (short crosslinking duration) *versus* 300 min (prolonged crosslinking duration, maximum structure formation). Lupin protein was crosslinked in at least 10 reaction tubes per sample type (i.e., 0 U TG/g protein, 10 U TG/g protein 60 min incubation, 10 U TG/g protein 300 min incubation). 1.25 g of the (crosslinked) lupin protein was weighed in one digestion test tube, representing one time point in the small intestinal phase (n=9 for 0, 10, 15, 20, 30, 45, 60, 90, and 120 min). To do that, aliquots were pooled from multiple tubes used during (crosslinked) protein preparation. This insured that the samples studied during digestion included variation that could have been introduced during the preparation stage. Thus, a kinetic digestion approach, adapted from previously reported work (Mukherjee, Duijsens, et al., 2026; Pälchen et al., 2021; Verkempinck et al., 2024), was used to monitor time-dependent structural changes as well as macronutrient hydrolysis kinetics. During the oral phase, 1.25 g of weighed sample was mixed with 1.0 mL simulated salivary fluid (SSF, pH 7.0) and 0.25 mL CaCl_2_ (7.5 mM), followed by brief vortexing. For the gastric phase, 2.0 mL simulated gastric fluid (SGF) and 0.05 mL CaCl_2_ (7.5 mM) were added to the oral bolus. The pH was adjusted to 3.0 with 2 M HCl, and Milli-Q water was added as described by Brodkorb et al. (2019). Pepsin (0.125 mL, 2000 U/mL digest) was then added, and samples were incubated at 37 °C for 120 min under end-over-end rotation (23 rpm). At the start of the intestinal phase, 2.125 mL simulated intestinal fluid (SIF, pH 7.0), 0.625 mL freshly prepared bile, and 0.4 mL CaCl_2_ (7.5 mM) were added. The pH was adjusted to 7.0 using 2 M NaOH to inactivate pepsin, and Milli-Q water was added to equalize volumes. An enzyme solution (1.25 mL) containing trypsin (100 U/mL digest) and chymotrypsin (25 U/mL digest) was added to obtain a final volume of 10 mL, followed by incubation at 37 °C for 120 min with end-over-end rotation (23 rpm). Digestion was terminated through protease inactivation through heating at 98 °C for 5 min, followed by cooling on ice. Samples were centrifuged at 2000 g for 5 min (Sigma 4-16KS, Sigma, Osterode am Harz, Germany), and the digestive supernatants were snap frozen and stored at −20 °C until analysis.

#### 2.8.2. Estimation of protein digestibility using OPA method

Protein hydrolysis during digestion was monitored by measuring the concentration of bioaccessible protein in the digestive supernatant using the OPA spectrophotometric assay, which quantifies α-amino groups (Nielsen et al., 2001) generated due to proteolysis. The procedure was adapted from established protocols (Mukherjee, Duijsens, et al., 2026; Pälchen et al., 2021). Total α-amino groups (*NH*_2(*total*)_) were determined after complete acid hydrolysis (24 h, 6 M HCl, 110 °C) of the undigested lupin protein isolate. In parallel, digestive supernatants were treated with 5 % Trichloroacetic acid (TCA) and centrifuged (Microfuge 22R (Beckman Coulter Inc, Indianapolis, IN, USA). TCA-soluble supernatant was hydrolysed (24 h, 6 M HCl, 110 °C) to obtain the TCA-soluble_hydrolysed_ fraction. This fraction consisted of free amino acids and represented the readily bioaccessible protein (Pälchen et al., 2021). Absorbance values were converted to L-serine equivalents using a standard curve (12.5–100 mg/mL), and protein digestion level as a function of time was expressed as the percentage of α-amino groups relative to those measured in the fully hydrolysed undigested sample using **equation 4**.

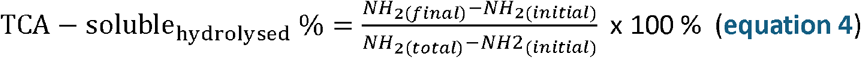

#### 2.8.3. Molecular size distribution of bioaccessible peptides: High-performance liquid chromatography size-exclusion chromatography (HPLC–SEC)

HPLC-SEC was used to characterise the molecular size distribution of bioaccessible peptides released during gastrointestinal digestion, following the approach described before (Duijsens, Pälchen, et al., 2023; Mukherjee, Duijsens, et al., 2026; Staes et al., 2026). Digestive supernatants were diluted in SDS buffer (0.05 M phosphate, 2 % SDS, pH 6.8), filtered (polyvinylidene fluoride membrane syringe filter, 0.45 µm), and analysed using an Agilent 1260 Infinity III LC system equipped with a diode array detector (Agilent Technologies, Santa Clara, CA, United States). Separation was performed on AdvancedBio SEC 300 Å and 130 Å columns (Agilent Technologies, Santa Clara, CA, United States) connected in series at 37 °C. The elution medium was SDS buffer (0.05 M phosphate, 2 % SDS, pH 6.8). Samples (10 µL) were eluted at 0.7 mL/min for 40 min, and absorbance was monitored at 205 nm. Data acquisition and processing were carried out using OpenLab Control Panel software (Agilent Technologies, Santa Clara, CA, United States). A calibration curve of molecular weight vs retention time was built using a protein standard mix (1–132.6 kDa,) (R^2^ = 0.97). Protein quantification was enabled by establishing a linear relationship between injected protein concentration and the area under the elution curve (AUC) using the mixed standard. The extent of proteolysis, expressed as bioaccessible content (BAC, %), was defined as the proportion of peptides ≤1 kDa and calculated using the AUCs of the corresponding SEC fractions (**equation 5**).

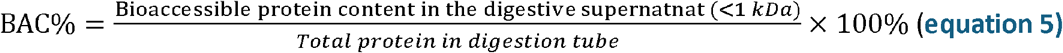

#### 2.8.4. Data analysis

Each digestion tube used for the small intestinal phase (n = 9; section 2.8.1) was treated as an independent measurement of the same system at a specific digestion time, following our multi-tube kinetic approach previously described in detail (Duijsens, Verkempinck, et al., 2023; Verkempinck et al., 2020). This approach allowed protein digestion kinetics for each sample to be characterised based on 9 separate observations. All data points obtained from sections 2.8.2 and 2.8.3 across the different digestion times were subsequently analysed by regression analysis; given the time-dependent behaviour of the hydrolysis data, a first-order fractional conversion model was used for the current work (**equation 6**).

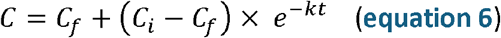

The digestive behaviour of the samples was then compared using the fitted kinetic curves and the corresponding model parameters. The parameters *C*_*f*_ (%) represent the estimated final extent of protein hydrolysis (expressed as TCA-soluble_hydrolysed_ % from section 2.8.2, or the bioaccessible protein % from section 2.8.3) at prolonged digestion times, and *k* (min^-1^), the estimated reaction rate constant. C_i_ (%) is the initial extent of protein hydrolysis and t is digestion time (min).

## 3. Results and discussion

### 3.1. Characterisation of the lupin protein isolate

The protein content of the lupin protein isolate was determined to be 85.1 ± 0.3 %. Furthermore, the isolate contained 9 ± 1 % lipids, 2.9 ± 0.1 % moisture, and 2.87 ± 0.02 % ash (wet basis). The protein had a solubility of 68.8 ± 1.5 %. Commercial lupin protein isolates are typically found to have a moderate to low solubility, often due to the drying step and/or heat treatment done to ensure microbiological safety (Porres et al., 2005). Furthermore, the solubility of the protein extract depends not only on the extraction process, but also on the variety of lupin seed (Igartúa et al., 2025). Fontanari et al., (2012) reported solubility values of 49.8 % for lab extracted protein from white lupin seed, whereas it ranged from 70.26 to 89.22 % for protein extracted from different varieties of blue lupin in the study of Bartkiene et al., (2018). Therefore, the blue lupin protein isolate used in the current study exhibited moderate solubility relative to values reported in literature.

DSC analysis revealed two distinct thermal transition peaks corresponding to α-and β-conglutin. α-conglutin had a denaturation enthalpy of 4.9 ± 0.3 J/g protein and a denaturation temperature of 94.6 ± 0.04 °C. β-conglutin had a denaturation enthalpy of 0.69 ± 0.04 J/g protein and a denaturation temperature of 87.8 ±0.2 °C (Wanniarachchi et al., 2026). Retention of endothermic peaks point toward retention of nativity or incomplete denaturation of the protein in the isolate. Comparing the denaturation enthalpy of protein extracts should be conducted with caution, as it is influenced by the extraction process of the protein, alongside the experimental conditions during the DSC analysis. Sirtori et al., (2010) reported denaturation enthalpy values of 0.70 and 0.85 J/g for β-conglutin and α-conglutin peak, respectively, in a blue lupin protein isolate prepared using air classification. Muranyi, Otto, et al. (2016) found that lupin protein extracted with micellar precipitation had a higher denaturation enthalpy compared with isoelectric precipitation (17.81 vs 10.44 J/g protein, respectively). The denaturation enthalpies and solubility measured in this study may indicate that the protein isolate likely contains a mixed population of native and partially unfolded proteins.

### 3.2. Development of viscoelastic networks in lupin proteins induced by TG

Gel formation of lupin protein dispersions (10 % (w/w)) was monitored using small amplitude oscillatory shear rheology to evaluate the gelation kinetics and viscoelastic behaviour of the system in response to different TG concentrations (0–10 U TG/g protein). The storage modulus (G’) was recorded throughout the incubation period at 40 °C for 5 h, followed by a thermal inactivation step at 70 °C for 10 min and subsequent cooling (**Fig. 1A**). For all treatments, G’ was consistently higher than G” (loss modulus; Supplementary Fig. 1), confirming the formation of a continuous network structure typical of gel systems. Similar trends have been previously reported for other legume proteins, such as pea (Djoullah et al., 2018), faba bean (Fan et al., 2024), or hybrid gels (Ye et al., 2026) undergoing TG-mediated crosslinking. Mukherjee et al. (2026) examined the effect of varying TG dosage and subsequent inactivation and cooling on the structure formation in pea protein. To the best of our knowledge, this is the first time a similar detailed rheological analysis of crosslinked lupin protein isolate has been conducted. In the control sample (0 U TG/g protein), minimal changes in G’ were seen during the incubation at 40 °C, whereas TG-treated samples showed a progressive rise in G’ during this phase (**Fig. 1A**). It was observed that TG-mediated crosslinking influenced both the gelation kinetics and the extent of rise of G’ during incubation. With increasing TG dosage, the rate of G’ development during incubation accelerated, and the final G’ at the end of incubation was higher, particularly for 1-10 U TG (**Fig. 1A, B**). This suggests that more extensive crosslinking and network densification occurred at higher TG levels (Vijayan et al., 2024). Across all systems, a further increase in G’ was observed during the inactivation step, which can be attributed to additional network formation driven by thermal gelation (**Fig 1B**). At the end of inactivation, the differences in G’ values across dosages were not statistically significant. After cooling, G’ reached a stable plateau, indicating that the gel structure had fully developed. While thermal treatment alone promoted some gelation, as indicated by the rise in G’ during the inactivation step, the final G’ of the control sample remained slightly lower than that of the TG-treated samples, although being not statistically significantly different. The rise in G⍰ observed during inactivation may partly reflect heat-induced modifications of protein. While exposure of hydrophobic patches and subsequent non-covalent associations represent a plausible mechanism consistent with previous reports (Ren et al., 2024), the current data do not permit direct determination of the interaction types. Thus, covalent crosslinking by TG leads to strong gel formation upon incubation (G⍰ > G⍰), but thermally induced gelation reduced differences among samples after cooling.

**figure 1.**
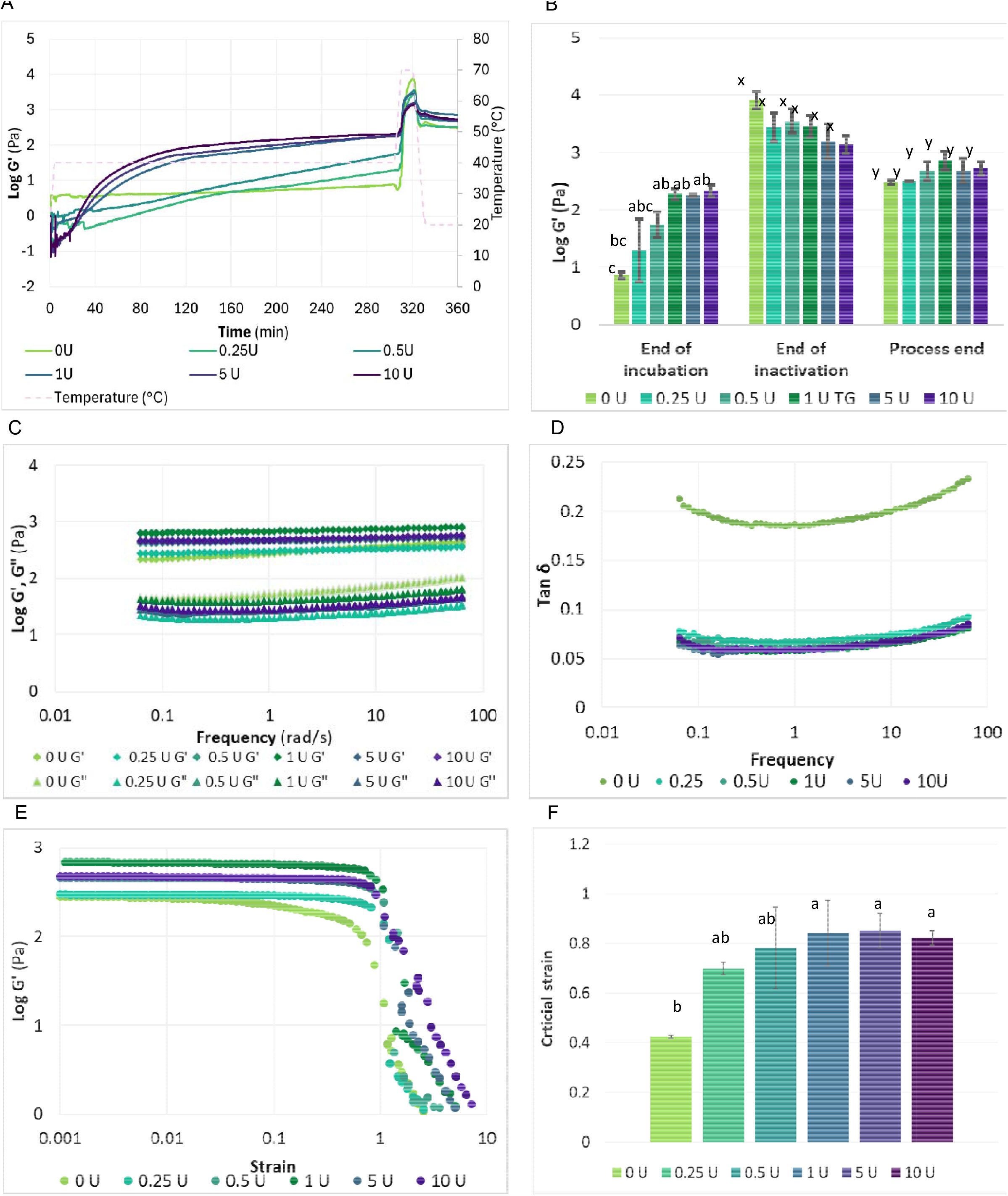
Rheological analysis of 10 % (w/w) lupin protein dispersions treated with 0, 0.25, 0.5, 1, 5, and 10 U of TG/g protein at 40°C and pH 7: me sweep showing the evolution of storage modulus (G⍰) over incubation and inactivation till process end. **(B)** G⍰ values during t e time swee and of incubation, inactivation and process end. **(C)** Evolution of storage modulus (G⍰) and loss modulus (G”) across frequency sweep **(D)** Tas G⍰/ G⍰) values across the frequency sweep. **(E)** Strain sweep depicting the linear viscoelastic region (LVR). **(F)** Critical strain for 0, 0. 5, 0.5, 1, 5, 1 G/g protein. All values shown are averages (measured in duplicate). The error bars indicate standard deviation; where not visible, error bars maller than the symbol size Different letters indicate statistically significant differences (p<0 05)

The viscoelastic properties of the final gels were further characterised using frequency and strain sweeps. The frequency sweep provides insight into gel stability by examining the mechanical spectra of the system across short and long time scales, represented by high and low frequencies, respectively (Moelants et al., 2014; Rao, 2014). As illustrated in **Fig. 1C**, G’ remained higher than G” across the entire frequency range for all samples, indicating that the gel network was stable and resistant to deformation. The weak frequency dependence of G’ is characteristic of covalently linked, well-structured protein gels (Bayod et al., 2008; Kavanagh & Ross-Murphy, 1998). The frequency dependence of G’ was quantitatively analysed using a power-law model (**Equation 2**). The model provided excellent fits to the data (R^2^ > 0.97), confirming its suitability for describing the rheological behaviour of the lupin gels. The consistency factors exhibited substantial variability across replicates, particularly at higher TG dosages (**Fig. 2A**). It is possible that higher TG dosages formed heterogenous networks as has been reported in literature (Rios et al., 2025; Wang, Jiao, et al., 2022). While the mean values suggest an increase up to 1 U followed by a plateau or slight decrease, these changes were not statistically significant. With increasing TG dosage, the exponent (b) decreased (0.11 ± 0.01, 0.04 ± 0.01, 0.032 ± 0.001 for 0, 0.25, and 10 U TG/g protein, respectively) (**Fig 2B**), indicating reduced frequency sensitivity (Ge et al., 2023). These changes reflect a progressive transition from weaker, thermally driven gels toward highly crosslinked, covalent networks as enzyme concentration increases. The parameter differences were statistically significant across TG dosages, underscoring the dose-dependent effect of TG on lupin protein network formation. Similarly, Yang et al., (2025) reported a transition from protein dispersion to frequency dependent, weak gels of TG crosslinked (2-14 U TG/g protein at 40 °C, 1 h) adzuki bean protein (10 % (w/v)) with electric field pre-treatment. While their system resulted in weak gel structures, our findings indicate the formation of more frequency independent, covalently crosslinked networks. Furthermore, tan δ values, representing the ratio of G” to G’, provided additional insight into the gel characteristics (Zhong & Daubert, 2013). In the current study, the tan δ of TG-treated samples remained rather constant across the frequency range (**Fig. 1D)**, consistent with stable, covalently crosslinked gels. The tan δ of the samples treated with 0 U TG/g protein, showed greater frequency dependence. Importantly, the average tan δ decreased with TG dosage: from 0.20 ± 0.01 for 0 U vs 0.07 ± 0.01 for 10 U TG/g protein. Values below 0.1 are typical for strong, elastic gels dominated by crosslinked structures (Bayod et al., 2008; Kavanagh & Ross-Murphy, 1998). Thus, it is clear that, although thermal gelation in the 0 U TG sample resulted in a relatively high G⍰ under small, linear deformations (**Fig. 1B**, process end), they show frequency dependence, typical of dense but weak networks gels (Hao & Weiss, 2013). The covalently crosslinked gels formed through action of TG, on the other hand, are strong, frequency-independent gels.

**Figure 2.**
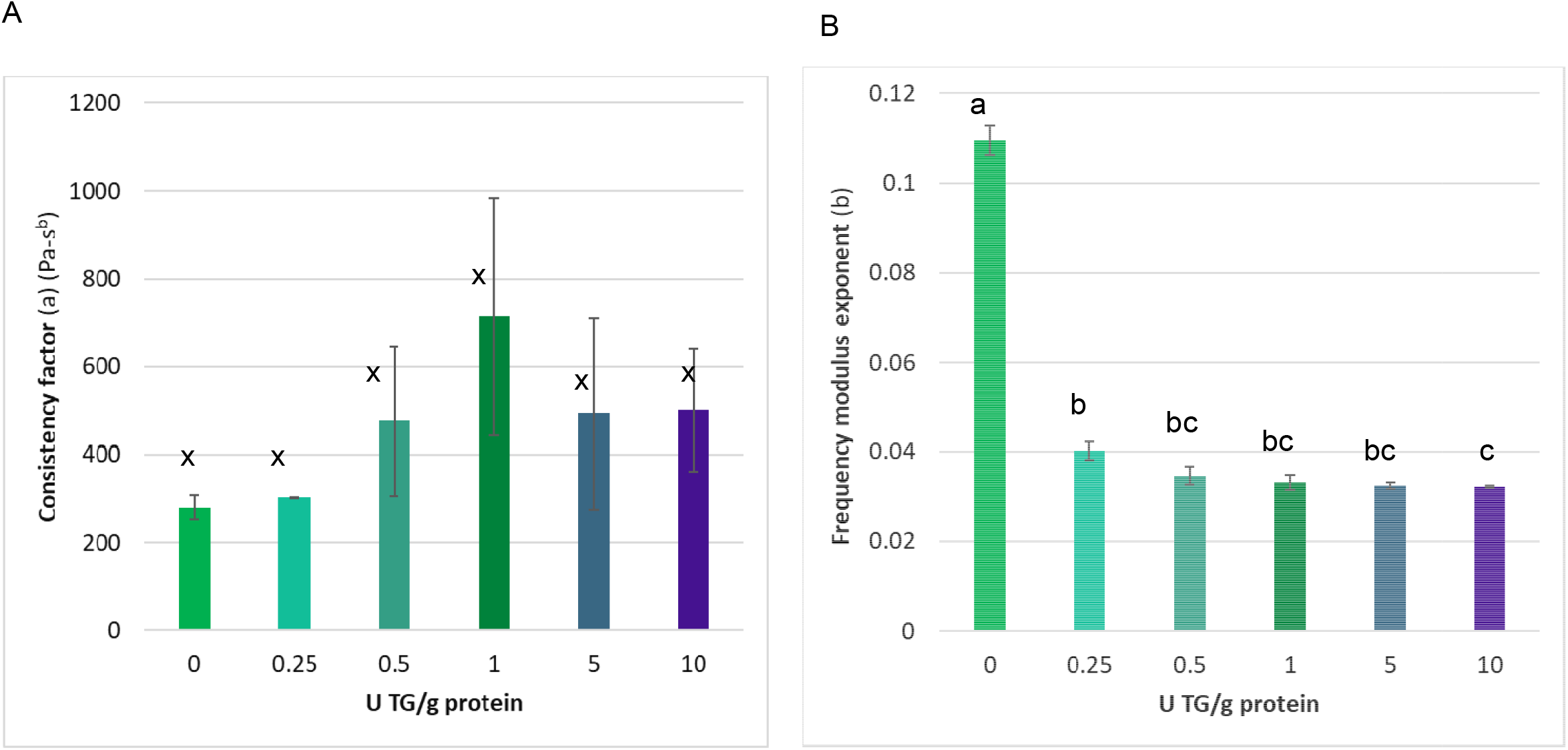
Frequency sweep parameters from the power law model (equation 1, section 2.5.4) of 10% (w/w) lupin protein dispersions treate with 0, 0.25, 0.5, 1, 5, 10 U of TG/g protein at 40 °C and pH 7: **(A)** consistency factor (a, Pa-s^b^). **(B)** Frequency modulus exponent (b). All value shown are averages (measured in duplicate). The errors bars indicate standard deviation, where not visible, error bars are smaller than the symbol size. Different letters indicate statistically significant differences (p<0.05).

Strain sweeps were performed to determine the LVR, defined as the range of strain values where G’ remains constant within ± 10 % of the preceding five measurements. As seen in **Fig. 1E**, TG treatment broadened the LVR, with the critical strain increasing from 0.42 ± 0.01 for 0 U TG to 0.82 ± 0.03 for 10 U TG (**Fig. 1F**). These results indicate that TG crosslinking enhanced the network’s ability to withstand deformation before structural breakdown. Similar findings were reported by Mukherjee et al. (2026) for TG-treated pea protein gels and by Masiá et al. (2023) for TG-treated pea protein emulsion gels developed using TG.

The results discussed above demonstrate that TG treatment of the lupin protein isolate yields a progressively stronger and more elastic gel network with increasing enzyme concentration. This dose-dependent effect, particularly observed during the incubation, reflects the role of TG in forming stable intermolecular bonds among the major lupin proteins, while also impacting overall gel structure.

### 3.3. Free ε-amino group content evolution as a function of enzyme dosage and incubation time

Since TG catalyses acyl-transfer reactions between the γ-carboxamide groups of glutamine residues (acting as acyl donors) and the free ε-amino groups of lysine residues (acyl acceptors), a progressive decrease in the concentration of free ε-amino groups is expected to occur as crosslinking proceeds. Monitoring the depletion of these ε-amino groups therefore provides an indirect measure of the extent of crosslinking. In this study, changes in free ε-amino group concentrations were quantified using the OPA assay, a common method for evaluating protein modifications during TG mediated crosslinking.

The experimental data (**Fig. 3**) revealed similar reaction profiles for all tested TG concentrations, characterized by a rapid initial decline in free ε-amino groups during the first hour of incubation (being less pronounced for 0.25 U TG), followed by a more gradual decrease as the reaction progressed. This behaviour suggests that the reaction follows pseudo-first-order kinetics, and the data were modelled using a fractional conversion approach (**Equation 3**) (Colle et al., 2010). The modelled curves indicated that the estimated rate constant (*k*) increased with higher TG dosage, rising from 0.007 ± 0.001 min^−1^ at 0.25 U TG/g protein to 0.017 ± 0.003 min^−1^ at 10 U TG/g protein (Table 1). Similarly, the final concentration of free ε-amino groups (NH_2final_) decreased as enzyme dosage increased, reaching 1.86 ± 0.03 mg/g sample for 0.25 U TG compared to 0.804 ± 0.086 mg/g sample for 10 U TG. This shows that with increasing TG dosage, a lower plateau in terms of the extent of crosslinking was reached faster.

**Figure 3.**
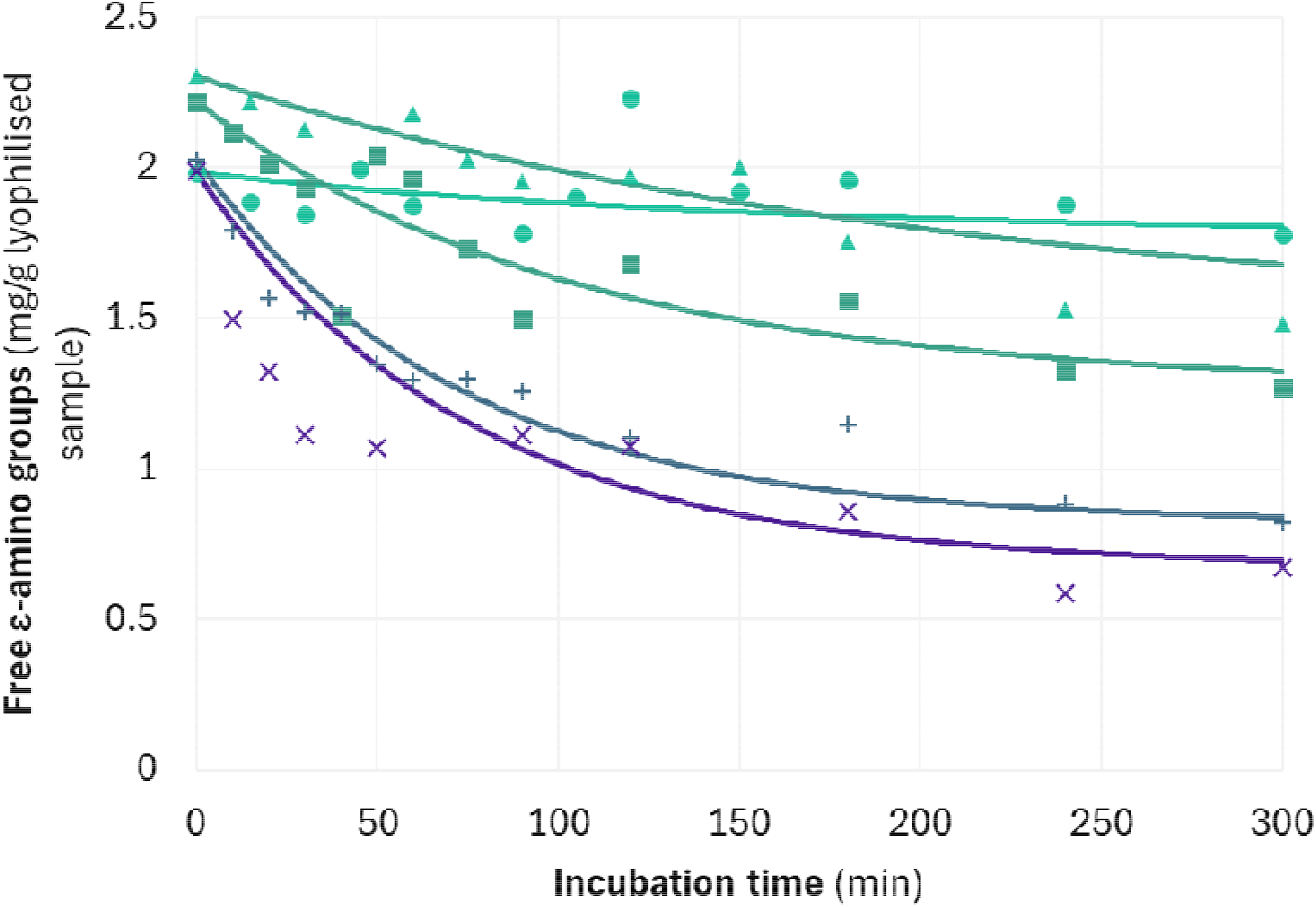
Change in ε-amino group groups in lupin protein (10 % (w/w)) crosslinked by TG (0.25, 0.5, 1, 5 and 10 U/g protein) at 40 °C, pH 7, as a function of incubation time. A fractional conversion model has been fitted to the kinetic data (equation 3, section 2.5.2). The symbols represent the experimental values and solid lines represent the modelled values.

**Table 1.** Parameter estimate of the rate constant, k (min^-1^) of lupin protein crosslinking (10 % w/v) by transglutaminase (0, 0.25, 0.5, 1, 5, and 10 U transglutaminase/g protein) incubated at 40 °C for 5 h. The parameter has been estimated from the decrease in free ε-amino groups due to the crosslinking reaction using the fractional conversion model (equation 3, section 2.5.2)

| TG dosage<br>(U/g protein) | $k$ ($\text{min}^{-1}$) | $\text{NH}_{2\text{final}}$ (mg/g<br>sample) | $R^2$ |
| --- | --- | --- | --- |
| 0.25 | $0.007 \pm 0.001$ | $1.86 \pm 0.03$ | 0.82 |
| 0.5 | $0.005 \pm 0.001$ | $1.02 \pm 0.2$ | 0.97 |
| 1 | $0.009 \pm 0.002$ | $1.3 \pm 0.2$ | 0.95 |
| 5 | $0.014 \pm 0.001$ | $0.95 \pm 0.05$ | 0.99 |
| 10 | $0.017 \pm 0.003$ | $0.804 \pm 0.086$ | 0.93 |

For the higher TG dosages, the depletion of free ε-amino groups reached a plateau within approximately two hours, suggesting that most accessible reactive sites had been utilised by that point. At lower TG concentrations, the decrease was more gradual and continued throughout the 300 min incubation. The deceleration of the reaction over time is consistent with progressive exhaustion of accessible reactive sites, as well as potential loss of enzyme activity during prolonged incubation. This progressive, dosage-dependent decline in free ε-amino groups has also been reported for other legume and cereal proteins, including pea (Mukherjee, Van Pee, et al., 2026), mung bean (Schlangen et al., 2023), and oat dough systems (Huang et al., 2010). The observed trend in lupin proteins aligns closely with the rheological results (section 3.2), where higher enzyme levels produced a faster and greater increase in G’ during incubation, indicating accelerated network formation. However, although different levels of ε-amino groups are reached at the end of incubation (and inactivation) at 300 min, the final G’ levels at the end of the process were not significantly different for the different TG dosages. Similar to our findings, where depletion of free ε-amino groups did not directly reflect gel strength, Schäfer et al. (2005) reported the highest ε-(γ-glutamyl)lysine isopeptide content after 120 min of incubation of lupin protein (13 % (w/v)) with 0.01 g TG/100 g protein at 40 °C. Nevertheless, only weak gels were formed, which may be attributed both to the comparatively low enzyme dosage used in their study and to differences in network architecture rather than total crosslink density alone. These results highlight that the number of crosslinks or extent of free ε-amino group depletion alone cannot predict gel network formation. Additionally, amino groups buried within a dense gel matrix may become inaccessible to the OPA reagent, leading to an underestimation of unreacted sites (Warakaulle et al., 2024).

Given these limitations, more detailed analytical techniques are needed to fully characterise TG mediated crosslinking and the distribution of crosslinks within lupin proteins. In this study, SDS-PAGE and proteome analyses were employed to determine which protein fractions, particularly the α-, β-, γ-, and δ-conglutins, participate most actively in TG-driven crosslinking. These results are presented and discussed in the following sections.

### 3.4. Molecular weight distribution of lupin proteins in relation to TG dosage: SDS-PAGE

The molecular weight distribution of the lupin protein isolate was evaluated using SDS-PAGE. As shown in **Fig. 4**, the lanes corresponding to the untreated lupin protein (NHNE) revealed distinct bands that align well with previously reported lupin protein subunits (J.Tang et al., 2025a). The major polypeptides identified included those around 70 kDa (high molecular weight (HMW) β-conglutins), around 50 kDa (acidic side chain of α-conglutins), 35 kDa (basic side chain of α-conglutins), 25 kDa (intermediate molecular weight (IMW) β-conglutins), around 30 kDa and around 17 kDa (likely corresponding to γ-conglutins), and <17 kDa (δ-conglutins) (Mazumder et al., 2021; Muranyi, Volke, et al., 2016; Tang et al., 2025b).

**Figure 4.**
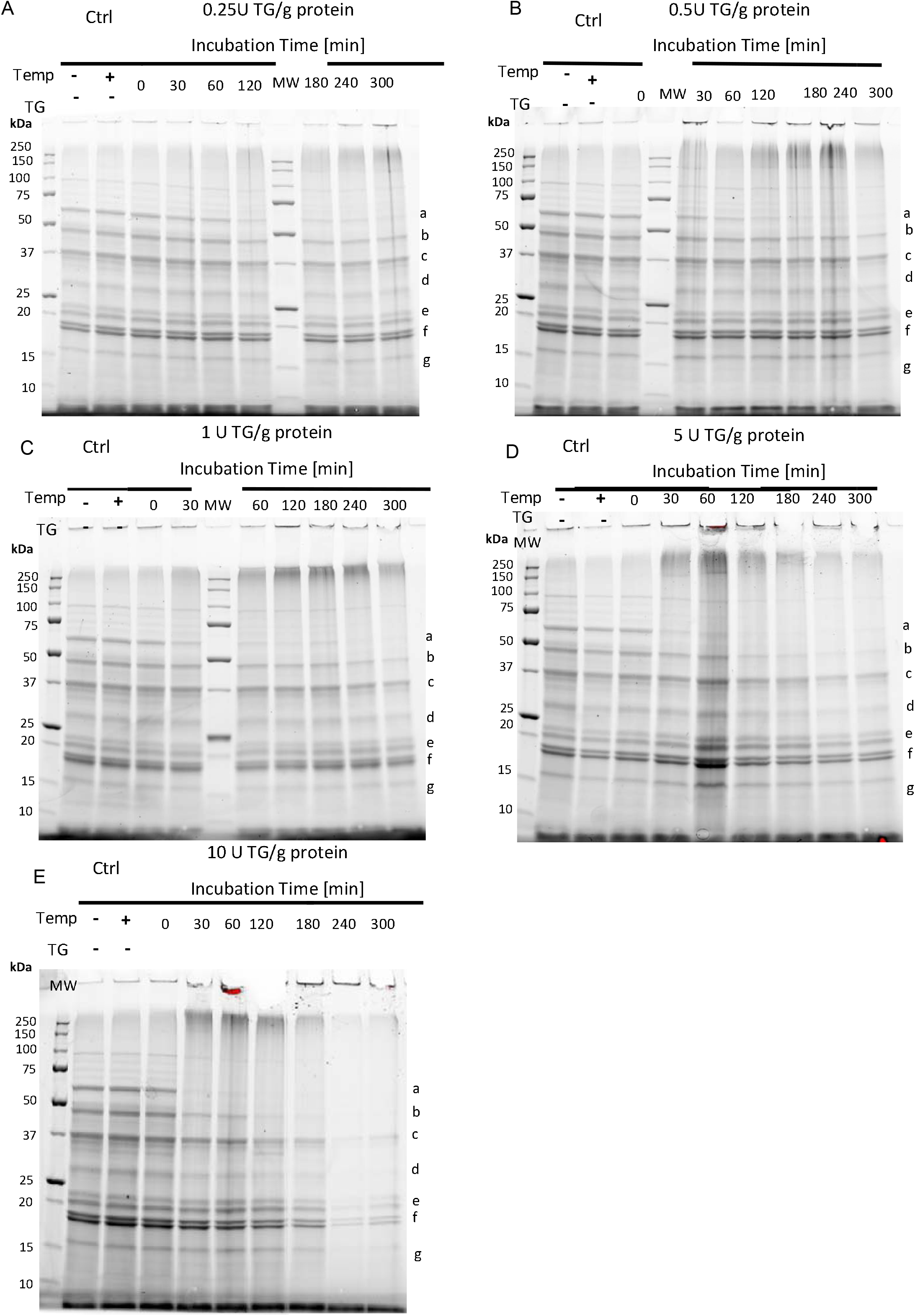
SDS-PAGE analysis of lupin protein (10% (w/w)) treated with **(A)** 0.25, **(B)** 0.5, **(C)** 1, **(D)** 5 and **(E)** 10 U of TG/g

Participation of the specific lupin proteins in the crosslinking reaction with TG was unravelled by SDS-PAGE analysis. The rationale was that TG-mediated crosslinking would lead to the formation of HMW aggregates, accompanied by a reduction in the intensity of specific bands as the proteins became covalently linked and migrated more slowly through the gel (Djoullah et al., 2018; Mukherjee, Van Pee, et al., 2026). Supporting this interpretation, new bands corresponding to HMW aggregates appeared at the top of the gel in TG-treated samples, while such bands were absent in non-enzyme controls (NHNE and HNE). This provides clear evidence that the observed aggregation was specifically driven by TG activity. A progressive decrease in band intensities was observed with increasing incubation time across all enzyme concentrations, indicating active participation of these proteins during crosslinking. It was observed that certain subunits were particularly reactive: α-(50 kDa) and β-conglutins (70 kDa) showed the most pronounced decrease, suggesting that these proteins were more accessible to TG. Additionally, the extent and rate of band disappearance were strongly dependent on TG dosage. At higher TG levels, the disappearance of α-and β-conglutin bands (70 kDa) occurred earlier and more extensively, indicating both faster reaction kinetics and a higher degree of crosslinking corroborating with the results from section 3.3. For samples treated with 10 U TG/g protein, the α-conglutin band (50 kDa) disappeared already after 60 min of incubation, whereas for 1 and 0.25 U TG/g protein, they started disappearing slowly only after 180 min. Similar observations were reported by Mukherjee et al. (2026) for pea protein, however, they observed a faster rate of disappearance of bands in general. They observed clear visual onset of disappearance of multiple bands at 10 U TG/g protein dosage already after 30 min. This may be due to structural differences of pea and lupin proteins or higher solubility of the pea protein used in their study which resulted in higher levels of crosslinking for the pea protein reported in their study. In contrast, lower molecular weight subunits such as δ-conglutins (17 kDa) exhibited limited changes in band intensity, implying lower reactivity or reduced accessibility to TG. The differential susceptibility of these subunits is likely related to their structural organisation and solubility characteristics (Yousefi & Abbasi, 2022). Previous studies on legume proteins have demonstrated that more hydrophilic and surface-exposed regions, such as those found in β-conglutins (and also to some extent in α-conglutins) (Nivala et al., 2017; Robinson et al., 2022), are more readily targeted by TG. In contrast, more hydrophobic or buried domains common in certain low molecular weight subunits are less accessible (Moreno & Clemente, 2008). Molecular-level characterisation of specific protein components, as presented in the following proteomics section, was conducted to further investigate these hypotheses.

### 3.5. Proteome-level estimation of TG crosslinking capacity

A key insight from our prior work on pea protein (Mukherjee, Van Pee, et al., 2026) was that TG-mediated crosslinking occurs between lysines and glutamines located preferentially in disordered regions. These regions provide the structural flexibility and solvent accessibility required for TG to access glutamine and lysine residues, particularly when supported by presence of particular residues (e.g., proximity to glutamic acid). Building on this mechanistic insight, a heuristic was developed to estimate TG-driven network-forming potential across plant storage proteins. The heuristic integrated two components: (i) the relative number of storage proteins capable of forming multiple TG-mediated connections (multi-connectors (>3 connections), chain-formers (2 connections), and terminators (1 connection)), and (ii) the balance of the reactive lysine and glutamine residues within disordered regions with glutamic acid motifs nearby. The product of these terms provided a composite heuristic used to rank species within the class of Magnoliopsida (complete list of 251,702,059 protein entries from 1,311,639 organisms in **Supplementary File 01**). Using this approach, known well working substrates for TG cross-linking emerged at the top positions: a.o. *Pisum sativum, Glycine soja, and Glycine max* ranked within the top 15 species, with blue lupin (*Lupinus angustifolius*) at position 24 (top 10 %). This positioned lupin as a promising candidate for TG-enabled structuring applications and was thus considered in the current study.

Furthermore, to evaluate whether the relationship between intrinsic disordered regions and TG-mediated crosslinking observed in pea protein still holds, the same data analysis pipeline (Mukherjee, Van Pee, et al., 2026) was applied to the blue lupin storage proteins. Firstly, the peptides exhibiting a negative log fold change (i.e., lower abundance after TG treatment), are peptides that were crosslinked by TG and thus modified. Hence, they were no more detected in the library search results as the Uniprot database does not contain peptides formed after post-translational modifications. Very limited number of peptides show a positive log fold change, which may be due to increased accessibility due to the thermal inactivation treatment enhancing the solubility of these peptides (Sharma et al., 2026). The results show a pronounced enrichment of crosslinked residues within disordered regions. The peptides exhibiting negative log fold change (**Fig. 5**), contained lysine and glutamine residues that were disproportionately associated with disordered regions. This trend was quantitatively robust: lysine residues are 6.37-fold (p= 1.23 × 10^-13^) more likely to be involved in crosslinking when located in disordered regions. Glutamine residues showed a 3.57-fold increase (8.84 × 10^-13^) under the same conditions. These enrichments indicate that structural accessibility, rather than simply residue presence, is a dominant determinant of TG reactivity. Similar results have been reported for other proteins. Nieuwenhuizen et al. (2003) reported that TG modified the glutamine and lysine residues only in the apo or partially unfolded states, while the native Ca^2+^-bound (holo) form at 30⍰°C is essentially unreactive in bovine α-lactalbumin. Spolaore et al. (2012) found that the main sites of modification of glutamine and lysine for apomyoglobin, apo-lactalbumin, and thermolysin fell within disordered regions. They further stated that the reactivity for lysine residues is generally less selective than for glutamine, as also noticed in the current study.

**Fig. 5.**
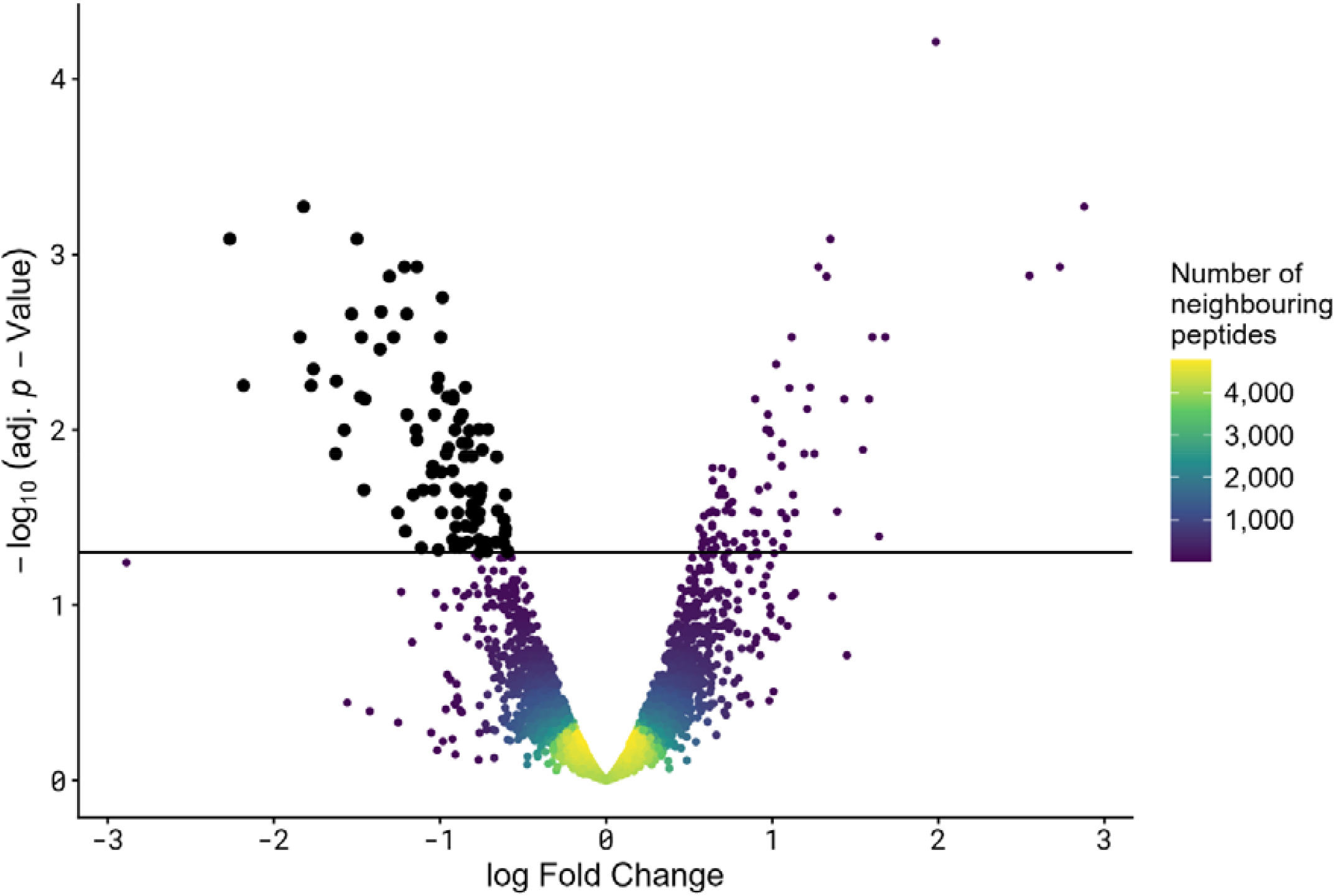
Volcano plot depicting the change in peptide abundance between TG crosslinked and untreated control from whole lupin protein lysate. Individual peptides are depicted. Black dots indicate over-representation of lower abundant peptides after TG treatment (negative fold change)

Taken together, these findings reinforce the hypothesis that intrinsically disordered regions act as primary reaction sites for TG-mediated crosslinking across plant protein systems. The consistency between pea and lupin strengthens confidence in using disorder-informed metrics, as predictive tools for identifying high-performing protein sources in TG-enabled structuring applications. To the best of our knowledge, this is the first time that in-depth proteomic approach has been employed to generate insight into domain specificity for TG catalysis in lupin protein.

### 3.6. Influence of TG mediated lupin protein crosslinking on *in vitro* protein digestibility

#### 3.6.1. TCA-soluble hydrolysed protein to estimate how the readily bioaccessible fraction changes as a function of *in vitro* digestion time

The *in vitro* protein digestion kinetics of the most distinct samples in terms of structure formation i.e., 0 U and 10 U TG/g protein (incubated for 60 and 300 min) were evaluated to understand the digestion outcomes of TG-mediated crosslinking of lupin protein. The extent of structure formation varied only slightly across the intermediate TG dosages (0.25-5 U TG/g protein), hence they were not considered for the digestibility study. The kinetic evolution of proteolysis was expressed in terms of percentage of TCA-soluble_hydrolysed_ protein (**Table 2, Fig. 6A**), representing the readily bioaccessible protein which can be absorbed by the brush border without further hydrolysis (Pälchen et al., 2021, 2022). The extent of gastrointestinal proteolysis, assessed by the OPA assay, showed a clear and consistent effect of TG-mediated crosslinking on lupin protein digestibility.

**Table 2.** Parameter estimates of the rate constant, k (min^-1^) and final level of TCA-soluble_hydrolysed_ protein C_*f*_ (%), during small intestinal proteolysis of lupin protein (10 % w/w) crosslinked by TG (0 and 10 U TG/g protein) incubated at 40 °C for 60 min and 300 min. The parameter has been estimated from the increase in free α-amino group due to the proteolysis reaction using the fractional conversion model (equation 6, section 2.8.4)

| TG dosage (U/g protein) | $k$ ($\text{min}^{-1}$) | $C_f$ (%) | $R^2$ |
| --- | --- | --- | --- |
| 0 | $0.16 \pm 0.02$ | $80.9 \pm 1.2$ | 0.98 |
| 10 (60 min incubation) | $0.09 \pm 0.01$ | $74.2 \pm 1.4$ | 0.96 |
| 10 (300 min incubation) | $0.09 \pm 0.03$ | $70.6 \pm 2.1$ | 0.96 |

**Table 2.**
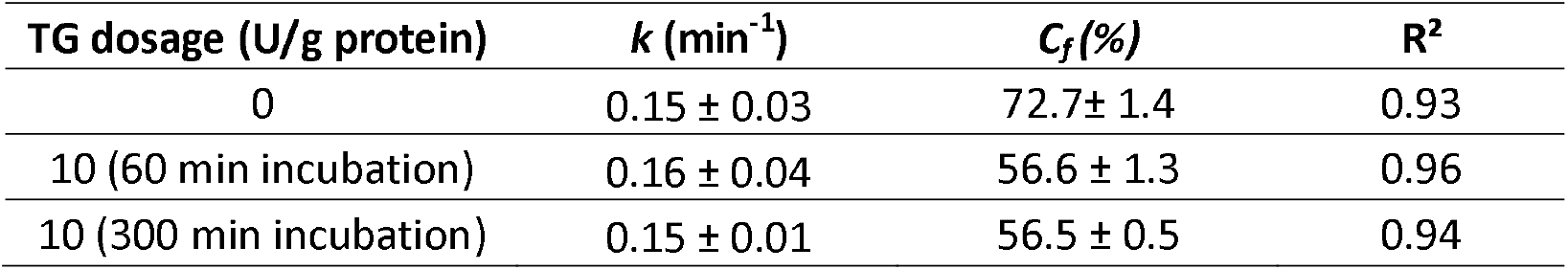
Parameter estimates of the rate constant, *k* (min^-1^) and final bioaccessible protein content C_*f*_ (%), during small intestinal proteolysis of lupin protein crosslinked (10 % w/w) by transglutaminase (0 and 10 U TG/g protein) incubated at 40 °C for 60 min and 300 min. The parameter has been estimated from the increase in peptides <1 kDa generated during gastrointestinal proteolysis, using the fractional conversion model (equation 6, section 2.8.4).

**Fig. 6.**
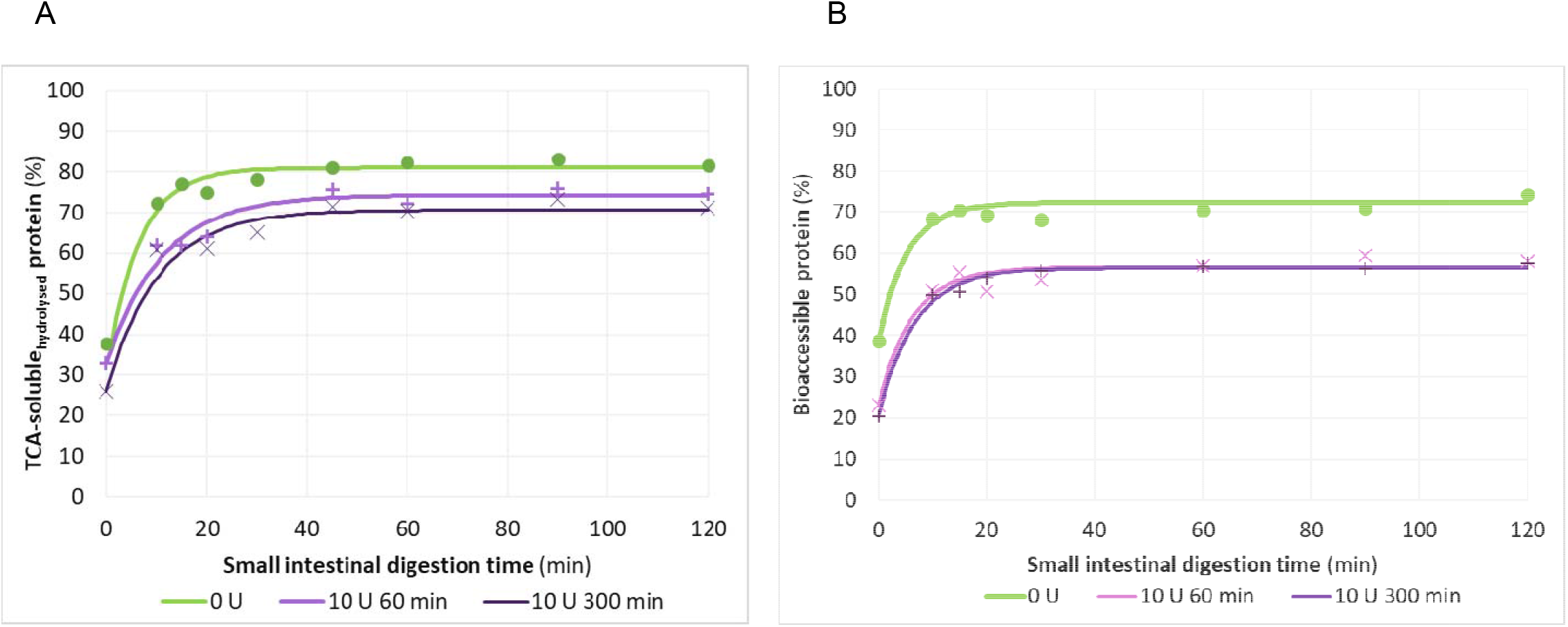
Protein digestibility during *in vitro* gastrointestinal digestion simulation of 10 % lupin protein (w/w) crosslinked with transglutaminase (TG) (0-10 U/g protein). **(A)** TCA -soluble protein_hydrolysed_ fraction **(B)** Percentage of bioaccessible protein (<1 kDa) of the digesta of lupin protein. Data has been modelled using fractional conversion model (equation 6, section .8.4). The symbols resent experimental values, lines represent modelled values.

All samples exhibited the characteristic digestion pattern of a rapid initial increase in TCA-soluble_hydrolysed_ protein during the early intestinal phase, followed by a gradual approach to a plateau. This trend reflects fast cleavage of accessible peptide bonds and subsequent limitation by substrate structure and protease accessibility. The untreated lupin protein (0 U TG) displayed the highest degree of hydrolysis at each digestion time point, maximally reaching approximately 80 % TCA-soluble_hydrolysed_ protein. This indicates that lupin protein, despite its globular nature, remained relatively accessible to pancreatic proteases once gastric pre-digestion was complete and intestinal conditions were established. The result aligns with previous reports that legume proteins such as faba bean, while partially resistant, can undergo proteolysis when not structurally reinforced by covalent crosslinking (Carbonaro et al., 2000). In contrast, TG-treated samples exhibited a reduced extent of hydrolysis, both in the 60 min TG-treated sample (around 75 %), and in the 300 min TG-treated sample (around 70 %). Thus TG-mediated crosslinking showed an effect on the rate and extent of hydrolysis. However, the effect was similar for the sample treated with 10 U TG/g protein for 60 and 300 min. This may be explained by the results from sections 3.2 and 3.3 (**Fig. 1A** and **Fig. 2**) where it was observed that prolonging the TG treatment did not cause a significant impact in terms of extent of crosslinking or structure generation. These findings strongly suggest that TG-induced crosslinking decreased the susceptibility to enzymatic hydrolysis. These findings are in agreement with previous reports of TG affecting the level of protein digestion in other plant protein systems (Mariniello et al., 2007; Romano et al., 2016b). However, contrasting results were presented by Wang et al. (2022), who found that for myofibrillar protein, limited crosslinking could increase protease accessibility. (Mukherjee, Duijsens, et al., 2026) reported a dose-dependent impact on the structural enhancement and the *in vitro* digestion kinetics of TG-crosslinked pea protein. They found that higher TG dosages (5-10 U TG/g protein) resulted in (s)lower digestion over time, with the strongest effects observed during the gastric phase. However, high digestion levels (70-80 % TCA-soluble_hydrolysed_ protein) were still reached later in the intestinal phase even for the extensively structured systems. It is also noteworthy that due to the more limited differences in the digestion behaviour observed, the range of targeted functionalities of food digestion design is narrower for the lupin protein under study compared to the pea protein. This underscores the protein and processing-specific nature of TG effects and the inherent structural rigidity of legume storage proteins, and thus prevents generalisation of literature results. Furthermore, it has been suggested in literature that the formation of ε-(γ-glutamyl)-lysine crosslinks likely decreased conformational flexibility and restricted access of digestive proteases to cleavage sites (Sabena et al., 2025). The digestion kinetics further support this interpretation. Although the initial rate of hydrolysis was relatively high for all samples, TG-crosslinked proteins showed slower hydrolysis already evident in the first minutes of intestinal digestion. This indicates that TG-mediated modification affected the accessibility of peptide bonds during the initial protease action.

Overall, these results demonstrate that TG treatment modulated the intestinal digestibility of lupin protein in a time-dependent manner when assessed by OPA. From a digestion perspective, the reduction in TCA-soluble_hydrolysed_ protein fraction suggests a potential decrease in protein bioaccessibility, particularly with extensive TG treatment (selected based on rheological and OPA assessments). However, as OPA primarily reflects the release of free ε-amino groups and small peptides, further analyses such as peptide size distribution is important to determine whether TG-induced crosslinking affects not only the extent but also the digestion quality of digestion products.

#### 3.6.2. Changes in protein size distribution and bioaccessibility during digestion of (non)crosslinked lupin protein: HPLC-SEC

HPLC-SEC analysis revealed digestion kinetics (Table 3, **Fig. 6B**) broadly similar to those observed by OPA, with a rapid formation of bioaccessible protein during the initial intestinal phase (0-30 min), followed by a stable plateau. The higher bioaccessible protein fraction observed for the 0 U TG/g protein sample indicates more extensive fragmentation into low-molecular-weight peptides. This behaviour is in line with earlier HPLC-SEC-based investigations, where less structurally constrained protein systems generated a larger proportion of peptides below the selected molecular weight cut-off during intestinal digestion (Duijsens, Pälchen, et al., 2023). In contrast, both 10 U TG/g protein (60 and 300 min) treatments exhibited substantially lower bioaccessible protein fractions, suggesting that a considerable portion of the protein remained either insoluble or were fragmented into peptides >1kDa threshold. Interestingly, extending the TG treatment from 60 to 300 min did not result in a lower bioaccessible fraction.

The rapid attainment of a plateau in all HPSEC-based proteolysis data is consistent with previous findings showing that the generation of low-molecular-weight peptides predominantly occurs early during intestinal digestion (Gwala et al., 2020; Pälchen et al., 2021). Continued intestinal proteolysis has been shown to yield peptides that either remain associated with insoluble fractions or do not further decrease in molecular size sufficiently to alter HPSEC-based protein bioaccessibility (Du et al., 2024; Duijsens, Pälchen, et al., 2023). Interestingly, the systematically higher digestion extents obtained by OPA compared to HPLC-SEC underline the different information provided by each method. The present results suggest that a fraction of peptides released during digestion contributes to the OPA signal without appearing in the HPSEC bioaccessible fraction. Previous studies have shown that OPA detects free α-amino groups originating from a heterogeneous mixture of peptides (Duijsens, Verkempinck, et al., 2023; Nielsen et al., 2001; Verkempinck et al., 2024). Such divergence between OPA and HPLC-SEC has previously been attributed to partial hydrolysis, or the formation of peptides that are soluble but yet still relatively large (e.g., >1 kDa). It has been further reported that differences in how individual amino acids and small peptides react with the OPA reagent, particularly those containing multiple primary amino groups such as asparagine, along with potential background interference, may lead to stronger OPA signals (Duijsens, Pälchen, et al., 2023; Mukherjee et al., 2026). Overall, the digestion behaviour observed here closely resembles patterns reported for structured plant protein systems (Picariello et al., 2023), reinforcing that protein accessibility plays a dominant role in determining both the extent and nature of digestion products. The combined use of OPA and HPSEC therefore provides complementary insight, allowing differentiation between peptide bond cleavage and the formation of low-molecular-weight, potentially absorbable peptides (Duijsens, Pälchen, et al., 2023).

## 4. Conclusion

This integrated, multiscale study, from macroscopic to molecular and digestion levels, provides a comprehensive understanding of TG activity on blue lupin proteins, linking structural functionality to digestion behaviour. Rheological analyses indicated the formation of a covalently linked protein network, while OPA measurements reflected changes in free ε-amino groups associated with TG-induced crosslinking. SDS-PAGE showed the progressive disappearance of specific protein bands (particularly of 70 kDa β-conglutin and the acidic and basic side chains of α-conglutin) as proteins became incorporated into HMW aggregates. Proteomics results indicated that lysine and glutamine residues located in disordered protein regions are more prone to crosslinking, likely due to their greater structural accessibility. The digestibility study revealed that lupin protein specific impact of TG on crosslinking and structure formation could be linked to the protein digestion outcome. Proteolysis was slowed and was lower than the non-crosslinked lupin protein. The effect was less outspoken than for pea protein (Mukherjee, Van Pee, et al., 2026). However, it is noteworthy that the study focused on a commercial lupin protein isolate. In future, different pre-treatment conditions such as a mild extraction process or high-pressure homogenisation may be explored to impart better solubility to the lupin protein and the consequences thereof on TG-mediated crosslinking. Amino acid analysis of the digesta can also be conducted to understand the impact of crosslinking on the release of essential amino acids such as lysine during gastrointestinal digestion.

## Funding Acknowledgements

A. Mukherjee is a doctoral researcher funded by Flanders Innovation and Entrepreneurship (VLAIO) in the context of ProFuNu project (HBC.2021.0546). D. Duijsens is a post-doctoral researcher funded by Research Foundation Flanders, Belgium (FWO - Grant no. 12A0225N). The authors acknowledge the financial support of Internal fund KU Leuven.

## Acknowledgements

We would like to acknowledge the FingerPrints Proteomics Facility at the University of Dundee, which is supported by the ‘Wellcome Trust Technology Platform’ award [097945/B/11/Z]. We would like to acknowledge Ilse De Pril and Annabel Meyfroot for their practical support.

