## Supplementary figure 01 for "Influence of transglutaminase mediated crosslinking on the structure-function-digestion properties of *Lupinus angustifolius* protein evaluated using a multiscale approach"

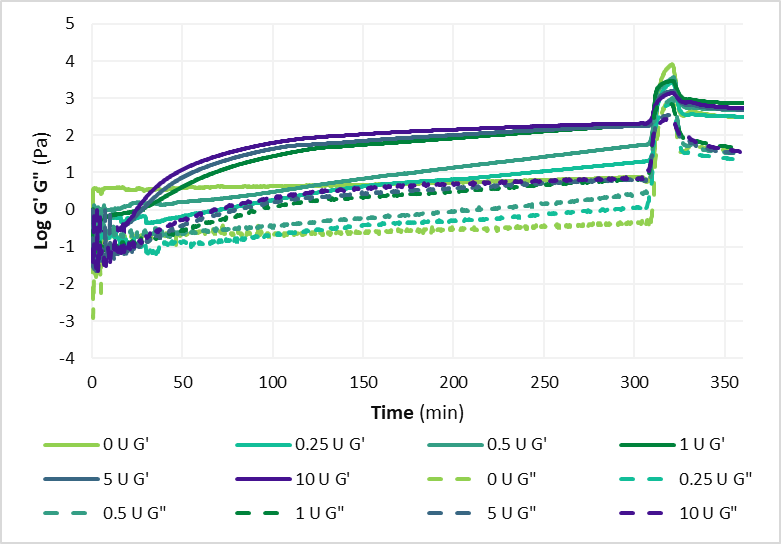

**Supplementary figure 1**. Rheological analysis of 10% (w/w) lupin protein dispersions treated with 0, 0.25, 0.5, 1, 5, and 10 U TG/g protein at 40 °C and pH 7: Time sweep showing the evolution of storage modulus (G′) (solid lines) and loss modulus (G”) (dashed lines) over incubation and inactivation till process end.
